# Effect of transcranial direct current stimulation (tDCS) on FRN and P2 during the performance of auditory monetary incentive delay task

**DOI:** 10.1101/2022.12.12.520071

**Authors:** Anastasia Grigoreva, Aleksei Gorin, Valery Klyuchnikov, Ivan Dutov, Anna Shestakova

## Abstract

Transcranial electrical stimulation (tES) serves as a powerful technique for assessing the causal role of specific brain regions in behavior, including decision-making. While tES studies investigating sensorimotor function yield unequivocal results, cognitive research reveals considerable ambiguity and diversity in stimulation-related effects. The consequences of transcranial direct current stimulation (tDCS) on cognitive functioning are not consistently predictable based on the current direction (cathodal or anodal), which limits its applicability in cognitive research.

In the present study, we further explored tES effect ambiguity in cognitive tasks by modulating medial frontal cortex (MFC) activity in an auditory monetary incentive delay (MID) task, where participants responded to acoustic cues encoding expected monetary losses using cathodal tDCS. We analyzed feedback-related negativity (FRN), reflecting prediction error processing when participants encountered losses compared to no losses during two subsequent MID task sessions, and exogenous P2 response to stimulus onset unrelated to anticipated monetary incentives. We anticipated an inhibitory cathodal tDCS effect on both P2 response and FRN.

Contrary to our expectations, we observed a facilitatory effect of cathodal tDCS on FRN, replicating our earlier results (Gorin et al., 2022). No effect of stimulation was observed on P2; however, tDCS influenced the learning effect of P2. The difference in P2 amplitude between the first and second sessions, observed in the sham group, was absent in the group that received cathodal tDCS. We offered the interpretation of the complex picture of tES effects during auditory MID performance in light of brain plasticity theory for P2 and reward-learning mechanisms for FRN. Importantly, our findings regarding the multidirectionality of tDCS effects on cognitive function challenge the utility of tES as a readily employable method for testing brain causality in highly complex neurocognitive events, such as decision-making.

## Introduction

The paramount methodological challenge in deciphering decision-making mechanisms pertains to the shift from correlational to causal models (Smith & Huettel, 2010). Conventional neuroimaging methodologies, encompassing electroencephalography (EEG) and functional magnetic resonance imaging (fMRI), possess the capacity to assess cerebral activity in conjunction with experimental variables. Nonetheless, to assert the causative involvement of a specific neural region in decision-making processes, it is essential to modulate brain functionality, while concurrently detecting the behavioral alterations engendered by manipulation. A multitude of such techniques has been extensively employed in animal studies, comprising optogenetics (Bernstein & Boyden, 2011), microstimulation (Hayden et al., 2008), and in vivo manipulation of neurotransmitter concentrations (Phillips et al., 2003). For the healthy human population, noninvasive methods of brain stimulation, namely tDCS are commonly used.

The transcranial direct current stimulation (tDCS) is a non-invasive technique that can directly affect cortical excitability, enabling its selective modulation (Been et al., 2007; Hanley et al., 2016; Boroda et al., 2020). According to somatic doctrine, anodal stimulation is hypothesized to enhance the excitability of the stimulated regions owing to cell depolarization (Batsikadze et al., 2013; Stagg et al., 2018). Cathodal stimulation is predicted to reduce excitability by hyperpolarizing the activated neurons (Batsikadze et al., 2013; Stagg et al., 2018; Nitsche et al., 2008). Still, effects of tDCS on brain function are multidirectional and often cannot be predicted precisely (Boroda et al., 2020; Stagg et al., 2018). Despite encouraging effects of transcranial electrical stimulation over the motor cortex (Nitsche & Paulus, 2000; 2001), applied to higher order neural networks tDCS effects are much less predictable (Jacobson et al., 2012; Kunzelmann et al., 2018; Reis et al., 2009; Wiethoff and Rothwell, 2014). Here, we investigated such a controversial tDCS effect on the medial frontal cortex (MFC) in a widely used reward-processing task called monetary incentive delay task or MID (Knutson et al., 2001) paradigm.

Reinhart and Woodman (2014) suggested earlier that learning processes can be causally controlled by tDCS. Namely, cathodal stimulation of MFC may decrease feedback-related negativity (FRN) and error-related negativity (ERN) compared to placebo due to MFC being responsible for the task performance monitoring, essential for reinforcement learning. Another tDCS study of To et al. (2018) shown, that using high definition anodal tDCS (HD-tDCS) led to significant rise in beta-band activity in dACC (anterior cingulate cortex), while cathodal HD-tDCS led to increased activation of theta frequency in dorsal and rostral ACC. Essentially, these study demonstrated the reliability of offline tDCS for dACC activity modulation. Here we used Monte-Silva et al. (2010) approach to spread offline effects of tDCS to all the time of MID task: 9+3+9 min (stimulation, rest, stimulation) of 1 mA current, allowing the effect to last up to 120 min.

The monetary incentive delay (MID) task is used to study neural mechanisms that provide stimulus-reinforcement association, allowing to examine prediction, anticipation, outcome processing and consumption in a variety of reward or punishment situations (Lutz and Widmer, 2014). Predicting and adapting behaviors based on experience (Noonan et al., 2012), individuals progressively change their internal perceptions of state-action values by measuring the difference between predicted results and external feedback (Sutton and Barto, 1998). The anterior prefrontal cortex (aPFC), the lateral orbitofrontal cortex (lOFC) and the ventromedial prefrontal cortex/medial orbitofrontal cortex (vmPFC/mOFC), are thought to greatly contribute to event value representation, reinforcement learning and decision making under uncertainty (Frank & Claus, 2006; Kringelback, 2005; Tsuchida et al., 2010).

Amplitude of the feedback-related negativity (FRN) correlates with these changes, in particular the sensitivity of FRN component to loss values (Gorin et al., 2020). Furthermore, it was found that, at the individual level, such changes in the amplitude of the MMN correlate with the sensitivity of FRN component to the magnitude of the monetary ouncome (Krugliakova et al., 2019; Gorin et al., 2020). FRN appears as the main candidate for neural correlate of reward prediction error (RPE) processing, observed in both the money loss and gain domains (Holroyd and Coles, 2002; Walsh and Anderson, 2012; Sambrook and Goslin, 2014). According to the reinforcement learning explanation, better learning and adaptation are driven by RPE (Hauser et al., 2014; Rouhani & Niv, 2021).

In our earlier work we demonstrated that FRN during MID task is sensitive to the magnitude of losses, dFRN as a difference between FRN in response to smaller and larger losses was negative in the low losses trials and positive in the high and distinct losses trials (Gorin et al., 2020). Our results indicate that in the context of losses, FRN is modulated by reference points and expectations. In a consecutive study (Gorin et al., 2022), we used cathodal tDCS to test whether inhibition of the medial prefrontal cortex, known to drive the generation of FRN in the context specific manner, be different depending on the context of the losses in the MID task: widely varying losses vs. low losses vs. high losses. In this study, we found that FRN amplitude in the loss prevention version of the MID task was enhanced by cathodal tDCS, although suppression was expected according to previous results obtained by Reinhart and Woodman (2014). Following the main assumptions of the reinforcement learning theory, we hypothesized that there is also a change in the perception of sound in terms of changes in event-related potentials (ERPs), and particularly there are such effects mediated by electrical stimulation in P2, which is by some researchers considered as the ‘biomarker of learning’ (Tremblay, 2014). Gorin et al. (2022), however, did not deserne the dynamic of FRN from the 1st to the 2nd session. Rather the researchers collapsed all trials from the first and the second day, seaking to the overall effect of TDCS.

We further hypothesized that if FRN was enhanced by cathodal stimulation, this change could be affected by 1) in the overnight dynamics of FRN and/or by the overnight dynamics of P200 (P2) event-related potential, a positive deflection with a typical peak latency of 150-250 ms induced by auditory stimuli (Ferreira-Santos et al., 2012). Studies examining different auditory training have shown that P2 amplitude increases with learning (Atienza et al., 2002; Hayes et al., 2003; Kuriki et al., 2007; Sheehan et al., 2005; Seppänen et al., 2012; Reinke et al., 2003; Kühnis et al., 2013) and represents the evaluation and classification of the stimulus (Tremblay et al., 2010). Wisniewski et al. (2020) observed a larger amplitude of auditory P2 accompanying trained sound detection rather than untrained in passive exposure of sound. Studies on the role of P2 in learning are not always able to establish a direct relationship between changes in P2 amplitude and the size of perceptual change, according to Sheehan et al., (2005), although P2 amplitude increase correlates with frequency modulation rate, reaction time (Tong et al., 2009), and increased performance in various tasks (Orduña et al., 2012). Notably, increased P2 amplitude is concomitant with improved processing of random sounds without attention: in a study by Seppänen et al. (2012), increased fast N1 and P2 plasticity reflected faster auditory perceptual learning in musicians.

Source of P200 is located mainly near the auditory cortex in the temporal lobe (Hari et al., 1990, Perrault and Picton, 1984). Among newer studies, Ross and Tremblay (2009) suggested that the center of P2 activity may be located in the anterior part of the auditory cortex, the lateral part of Heschl’s gyrus. In study by Crowley and Colrain (2004), the generation of P2 is dedicated to planum temporale, Brodmann’s area 22, and auditory association cortices. Continuing the assumption of finding the source of P2 in the temporal lobe, there are reports that tissue loss in the frontal lobe also correlates with P2 amplitude in the attention task, but if the source of P2 generation is in the temporal cortex, other brain regions, such as the frontal cortex, may modulate this component (McCarley et al., 1989, 1991). Other reports suggest that the inferior parietal lobe can also modulate the P2 signal (Knight et al., 1988). In addition, this component may also take its origin from sources outside the cortex (Ferreira-Santos et al., 2012; Jacobson, 1994). MEG and Intracranial EEG data also suggest that P200 has different sources: the temporal plane and the auditory association complex in area 22 (Godey et al., 2001). Thus, most studies suggest that P2 originates from associative auditory temporal regions, although there may be non-temporal sources active in the same time window.

To investigate cathodal tDCS effects on reinforcement learning-related cortical plasticity evoked by the two-day performance of MID task, a sham-controlled study was conducted. Montage as in Reinhart and Woodman (2014) over the medial frontal cortex was used.

The present study is designed to study to what extent the facilitatory effect of cathodal tDCS on FRN could be explained by the tES modulation of auditory N1-P2 responses, which by some researchers considered as the ‘biomarker of learning’ (Tremblay, 2014). These two components appear to reflect different processing associated with the repeated performance of the MID task and possibly activate different networks. We used tDCS to trace the plasticity of the two mechanisms and how they intertwined in the MID task performance.

## Materials and methods

### Participants

30 healthy right-handed participants (18 women aged 23+-2 years) were invited to take part in the study. Half of the participants formed a cathodal stimulation group (Reinhart and Woodman (2014) montage), while the other half received sham (placebo stimulation) just before the MID task. The study protocol was approved by the local ethics committee. All participants gave informed written consent before entering the study.

### Procedure

The experiment consisted of 2 day-by-day sessions, including stimulation (sham or cathodal) for control and experimental group respectively, and the MID task (Figure 1). Participants got a quantity of money in the amount of 4’000 rubles before the MID session. They were told that throughout the game session, they may lose some of the money, and that their reward for participating would be based on the remaining amount. A modified version of the MID task was used for the experiment as in Gorin et al. (2020).

**Figure 1.**
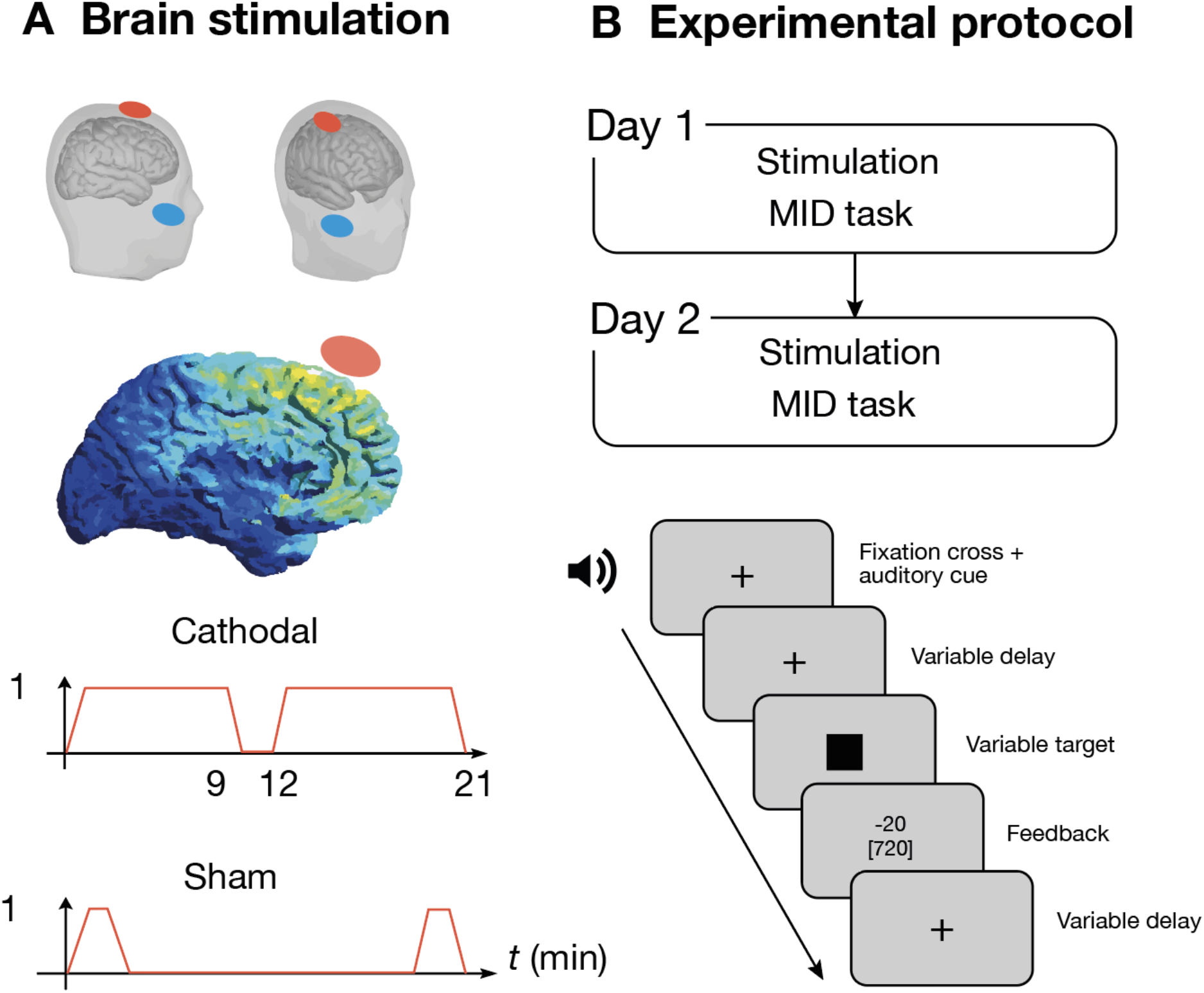
Study design. (**A**) Brain stimulation montage over MFC. Model of current distribution in the stimulating electrode assembly used (performed in SimNIBS), stimulation protocol. Cathodal tDCS is expected to deliver the current to MFC and ACC. (**B**) Study and MID task structure.

Six auditory stimuli (325, 381, 440, 502, 568, and 637 Hz) constituted three pairs of stimulus signals that predicted low and high losses: -1 or -2, -50 or -51, -1 or -50 rubles (low losses – LL, high losses – HL, distinct losses – DL, respectively). Thus, the 1-ruble difference between the outcomes in the LL and HL settings did not matter considering the beginning sum of 4,000 rubles, while the 49-ruble difference between the outcomes in the DL context mattered. Participants were therefore more driven to recognize signals in the DL setting to enhance monetary outcomes.

Each pair of auditory money signals was randomly presented in mini blocks of 50 trials. In total, each session of the MID task consisted of six mini blocks, so that each of the three context types appeared twice during the experiment. The sample size was too small for full randomization; therefore, the acoustic match between stimulus and outcome using six main combinations was counterbalanced. Thus, the signal-result match between lower/higher frequency and lower/higher outcome values was counterbalanced among participants.

tDCS was performed using a StarStim electrostimulator and two conductive rubber electrodes. Elliptical (active) and rectangular (referent) electrodes were placed in saline-soaked synthetic sponges (active electrode, 19.25 cm^2^; reference electrode, 52 cm^2^) and fixed in proper positions using a cap and bandage. The electrode positions were like in the work of Reinhart and Woodman (2014). Cathodal tDCS was expected to deliver the current to MFC and ACC. The active electrode was positioned in the FCz position according to the 10-10 system, and the referent was put on the cheek, left or right (equally in each group, reproducing approach used in Reinhart and Woodman, 2014). The current strength was 1 mA; the duration of the effect was extended using a customized stimulation pattern: 9 minutes of stimulation, followed by 3 minutes of rest, and then another 9 minutes of stimulation, which ensured (Monte-Silva et al., 2010) that the post-stimulation effect would last throughout the MID task session.

During the game, EEG and EOG were recorded. EEG recording was acquired using the BrainProducts ActiChamp system with 60 active electrodes arranged according to an extended version of the 10-20 system. The signal was abstracted to the average signal from a pair of electrodes on the mastoid processes. An electrooculogram (EOG) was recorded using electrodes placed under the right eye and on the left cheekbone. The ground electrode was placed in the Fpz position. The impedance of the electrodes was maintained below 5 kOhm before recording.

### Data analysis

#### EEG preprocessing and ERP analysis

Data analysis was conducted using MatLab Brainstorm (Tadel et al., 2004) and R (R Core Team, 2022), figures were produced using the package ggplot2 (Wickham, 2009) and ggstatsplot (Patil, 2021). Raw data was checked manually and then eradicated of artifacts, labeled relative to auditory stimuli (coded loss amount), context of playing and trial; averaged within each group, computing both subject and grand averages. EEG data was bandpass filtered in the range from 1 to 40 Hz, eye movement artifacts have been removed using independent component analysis according to their topography and correlation with the EOG. Noisy segments were marked as bad in the recordings during preprocessing and were excluded from further analysis.

After the preprocessing, epochs relevant to each stimulus in range of -200 and 800 ms with preprocessing of removing DC onset and selecting baseline definition in time range of -100.0 and - 2.0 ms were imported to the database. Time window of 100-300 ms relative to the auditory stimulus, as in a previously published study (Krugliakova et al., 2018) was analyzed. The obtained periods were averaged within experimental (cathodal) and control (sham) groups for each day, across trials, game contexts, and loss sizes for all electrode positions.

#### Statistical analysis

To assess the significance of the observed differences in ERPs between the two groups and within each group, permutation tests were employed. Paired permutation tests were conducted for both control and experimental groups across all days, contexts and losses. For example, a pairwise permutation test was performed between day *1* and day *2* data for the *cathodal* group with 1-ruble losses (*lesser amount*) in the *DL* context. Independent sample permutation tests were also performed between *sham* and *cathodal* stimulation groups in general, and within the same context, loss size, and day. Results of permutation tests were adjusted for multiple comparisons using the FDR (false discovery rate). *When the data was extracted for further investigations, t-tests were partially replicated for only the Cz channel*.

Subsequently, repeated-measures analysis of variance (RM ANOVA) was conducted with stimulation (cathodal, sham) as a between-subject factor, stimulus value (lesser or greater amount), context (LL, HL, DL), day (1, 2) as within-subject factors for P2 amplitude. The threshold for statistical significance was set at an adjusted p-value of 0.05, the results were Bonferroni corrected.

FRN response was calculated as a difference between no-losses and losses trials at the time of the feedback given to the participant. FRN modulation was earlier demonstrated (Gorin et al., 2022) with pairwise permutation tests between the stimulation and placebo groups in pairs for each type of monetary loss, focusing on a time window of 200-300 ms. The effect of stimulation on the FRN amplitude depending on the conditions and context of the experimental task was assessed by rm ANOVA with the factors *stimulation* (cathodal or placebo), *context* (LL, HL, DL), and *loss magnitude* (lesser or greater).

Here, in order to monitor the effect of the MID training, we also split FRN values in two groups based on the *day* of session and fit the extracted amplitude values (200-300 ms) to rm ANOVA.

#### EEG induced activity analysis

The study also assessed oscillatory responses to sound cues in the monetary game. Raw data were re-referenced to averaged potential and segmented into epochs of -1000 to 2000 ms. Time-frequency transformation was applied using the Morlet wavelet function, focusing on the 4-20 Hz frequency range, encompassing alpha, beta, and theta bands. The phase-linked component was removed, and data were standardized into an event-related synchronization-desynchronization view. Pairwise permutation tests were utilized to compare frequency-time dynamics between experimental sessions.

## Results

### Effect of cathodal tDCS on N1 and P2

We recorded EEG before and after two day-by-day MID task sessions. Fig. 3 shows the ERP curves for the two groups, averaged over the stimulation conditions, the size of the encoded loss, and the day.

**Figure 3.**
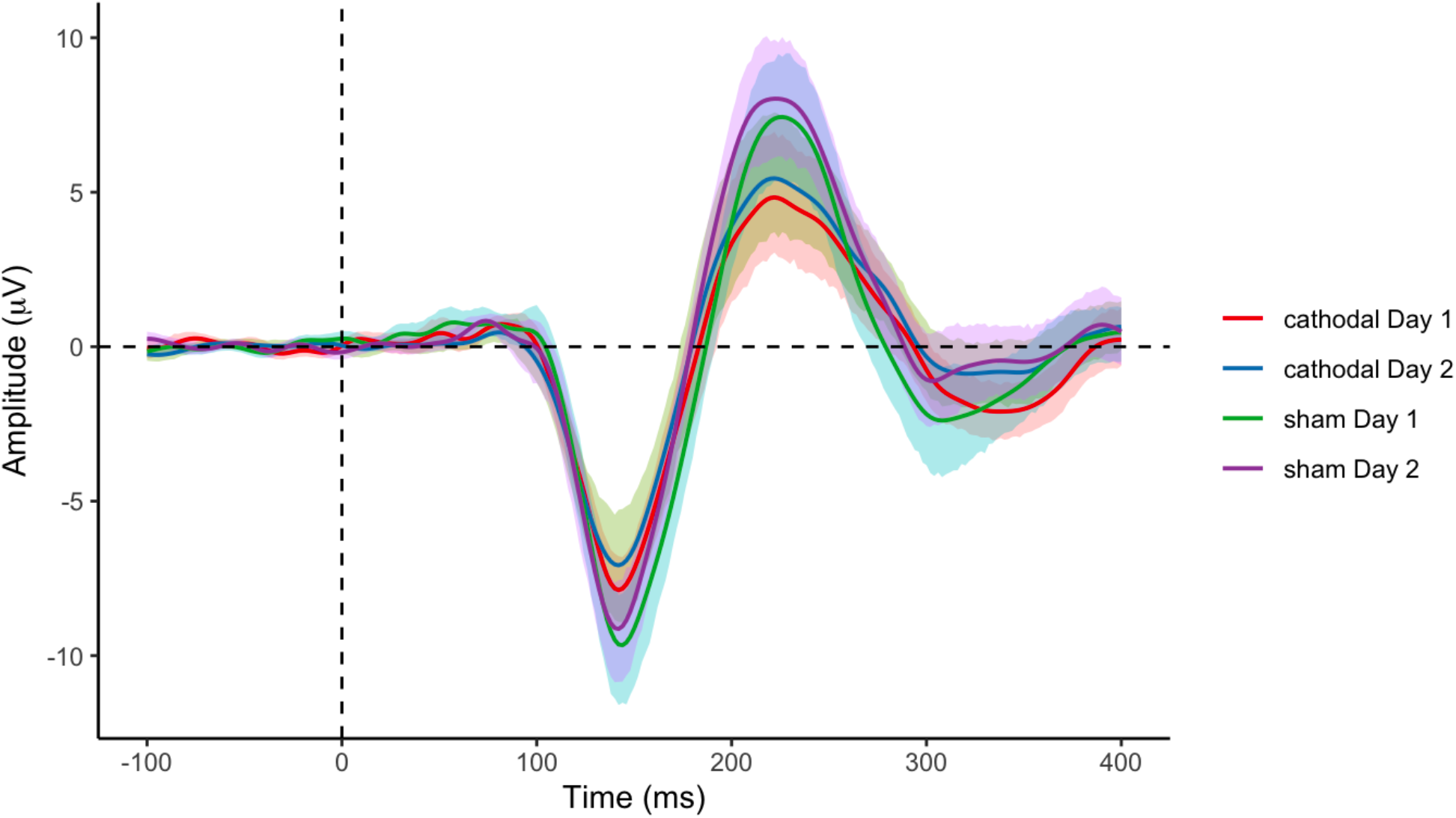
General average for experimental (cathodal stimulation) and control (sham) groups for 1^st^ and 2^nd^ day with confidence intervals (95%). Cz electrode. P2 amplitude is significantly higher in the control group, both on the first and the second day (p<0.05).

Noticeably, P2 amplitude is larger in the control group that received sham stimulation compared to the cathodal stimulation group, and particularly for the second day. Then we performed an unpaired permutation test comparing ERPs on each channel in cathodal stimulation condition vs sham condition on the first day. A series of permutation tests for the **N1** amplitude revealed **no significant differences** in the time range of the studied component in all conditions. Besides, permutation tests revealed a significant effect of the group on the amplitude of the **P2** component: in the *sham* condition, the P2 component showed enhanced day-to-day magnitude, while *no P2 amplitude increase in stimulation group was observed while significant P2 amplitude increase in sham group*.

Next, we compared the difference ERPs (day two – day one) in the control group pairwise for each type of money loss, focusing on a time window of 100-300 ms. Results were adjusted for multiple comparisons using the FDR (false discovery rate) method. A pairwise permutation test showed significant differences (Fig. 4) at the 95% confidence level of 2 vs. 1 day in all game contexts in the following time intervals: for DL context -1 R 180-212 ms – smaller amount; for DL context -50 R 174-204 ms and 258-284 ms – larger amount; for HL context -50 R 166-212 ms and 260-292 ms – smaller amount; for HL context -51 R 170-212 ms and 272-278 ms – larger amount; for LL context -1 RR 188-212 ms – smaller amount; for LL context -2 RR 170-214 ms and 272-284 ms – larger amount.

**Figure 4.**
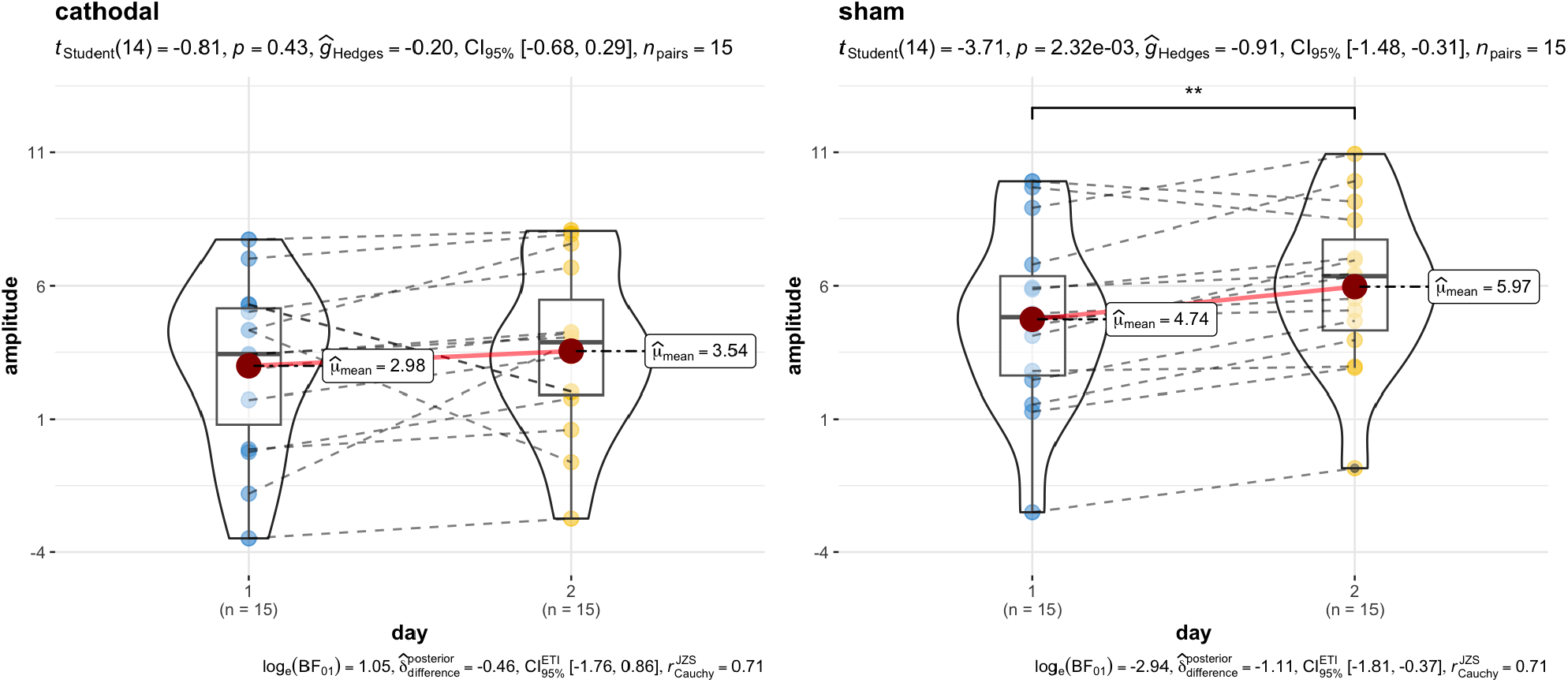
Results of t-test of P2 amplitude (Cz electrode) by the factor Day, grouped by type of stimulation. Amplitude growth is present only in the sham stimulation group, the effect size in control group is large (*g*=-0.91). Bayes factor also confirms positive evidence for H1 in the sham group. In the cathodal stimulation group, the growth is noticeably smaller, and the growth is not significant.

**Figure 5.**
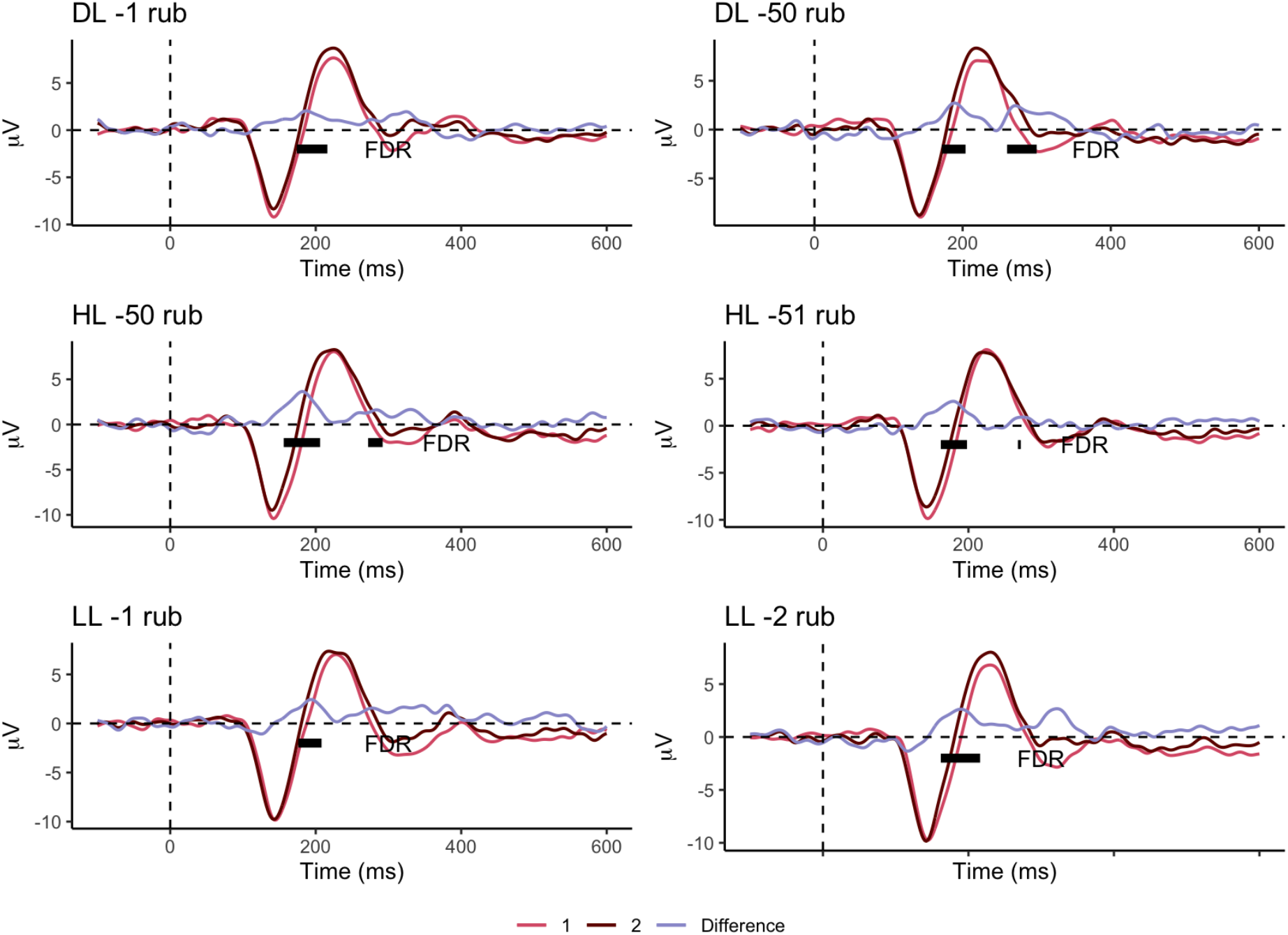
ERP curves for sham group according to Day, Cz electrode. The difference between the ERPs is pronounced in all contexts of play presented (p<0.05, FDR correction).

Since significant differences were found for all loss contexts played (p<0.05), we extracted P2 values for the difference between 2^nd^ and 1^st^ day and got repeated measures analysis of variance. To test the significance of the P2 differences, we averaged the signal observed on the Cz channel in the 180-260 ms time window (Magnée et al., 2008) and performed RM ANOVA with the factors: day, context, and loss size (Table 2 and Figure 6). The analysis of variance of the amplitude of the P2 component revealed the significance of the factor Day (F (1, 13) = 11,182, p = 0,002, η_p_^2^ = 0,493645).

**Table 2.** Repeated Measures Analysis of Variance with Effect Sizes and Powers for P2 amplitudes

|  | SS | DF | MS | F | P | Partial eta-squared | Observed power |
| --- | --- | --- | --- | --- | --- | --- | --- |
| Day | 69,02 | 1 | 69,02 | 13,65 | ,002 | **0,494\*** | **0,929\*** |
| Context | 15,28 | 2 | 7,64 | 2,01 | 0,153 | 0,126 | 0,380 |
| Value | 3,31 | 1 | 3,31 | 1,24 | 0,284 | 0,081 | 0,180 |
| Day*context | 1,13 | 2 | 0,56 | 0,27 | 0,762 | 0,019 | 0,089 |
| Day*value | 0,11 | 1 | 0,11 | 0,09 | 0,764 | 0,007 | 0,059 |
| Context*value | 1,35 | 2 | 0,68 | 0,52 | 0,599 | 0,036 | 0,127 |
| Day*context*value | 2,55 | 2 | 1,28 | 0,7 | 0,505 | 0,048 | 0,156 |

Notes. \* $p < 0.05$

### FRN amplitude is larger than expected

To test the differences in FRN, we averaged the signal observed on the Cz electrode over a time window of 200-300 ms and used repeated measures analysis of variance with the factors *Stimulation* (cathodal or sham) and *Day* (1 or 2). The variance analysis revealed a significant effect of *Stimulation* (F (1, 29) = 5.75, p = 0.023, η_p_^2^ = 0.16), where cathodal stimulation resulted in a significantly more pronounced FRN signal than sham (-2.6 μV and -0.91 μV, respectively; see *Figure 7*). The influence of the Day factor was insignificant, as was their interaction (p > 0.13).

**Figure 7.**
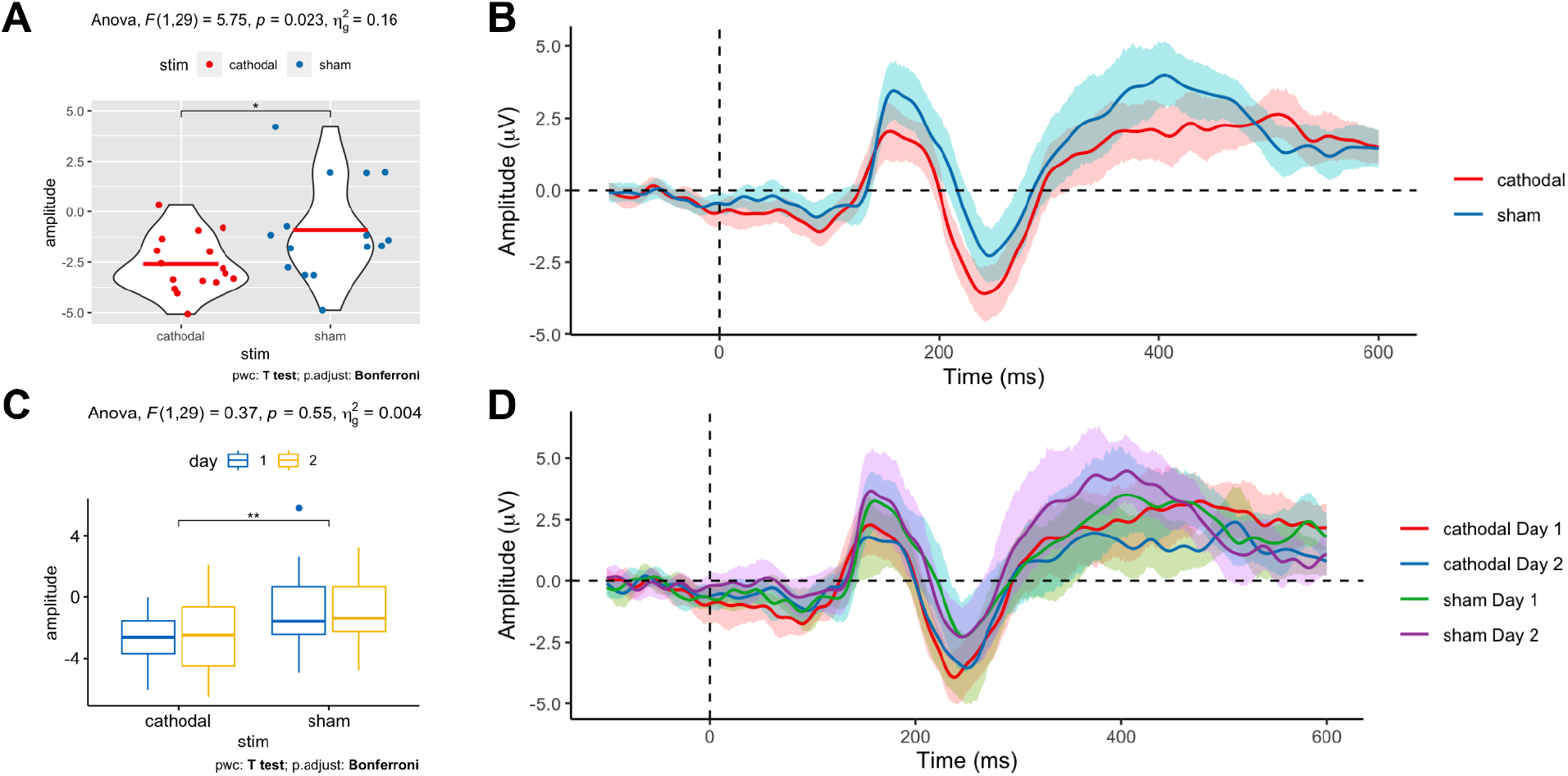
(**A**) Results of analysis of variance by the factor *Stimulation* between sham and cathodal groups. The FRN magnitude in the cathodic stimulation group was significantly higher than in the control group, p = 0.02. (**B**) EP curves for the two groups, averaged under *Stimulation* condition, Cz electrode. (**C**) RM ANOVA for interaction of *Day* and *Stimulation* factors, a significant effect is presented only for *Stimulation* factor. (**D**) EP curves for two groups averaged under *Stimulation* and *Day* condition, Cz electrode.

## Summary and Discussion

In this study employing two sessions of MID task, we focused on the effect of cathodal tDCS on the brain plasticity manifested in the P2 component changes from one day of MID task performance to another. Our previous research revealed that the FRN response in the MID task was significantly higher in the group which obtained stimulation than in the control group (Gorin et al., 2022) when the trials from both days were collapsed together. FRN seems to represent the computation of RPE, indicating outcome and prediction inconsistency (Holroyd et al., 2004; Sambrook and Goslin, 2014; Walsh and Anderson, 2012). According to the reinforcement learning theory, larger RPE provides better learning (Rouhani & Niv, 2021). According to Kiat and colleagues (2016), P2 also plays a role in the neural coding of outcome predictability and is sensitive to perceptual learning, memory, and training (Habibi et al., 2014). These two components appear to reflect different processing associated with the repeated performance of the MID task and possibly activate different networks. We used tDCS to trace the plasticity of the two mechanisms and how they intertwined in the MID task performance.

We observed no significant change in N1 depending on day or simulation. In the group that received the placebo stimulation, we observed an increase in the amplitude of the P2, which many studies (Atienza et al., 2002; Sheehan et al., 2005; Tremblay et al., 2001; Reinke et al., 2003; Hayes et al., 2003) attributes to plastic changes in the brain. On the contrary, in the group that received cathodal stimulation, we found no significant difference in amplitude between the experiment’s first and second days. This observation is correct for each of the six monetary loss conditions: -1 and -2, -50 and -51, -1 and -50 rubles. Since the P2 amplitude on the first day is significantly higher in the control group, we assume that cathodal stimulation suppressed the P2 amplitude for both days of the experiment. The oscillatory data is consistent with the observed P2 inhibition. Thus, we can assume that cathodal stimulation influences the changes in the electrical activity of the brain, at least by slowing down their appearance (Holroyd et.al., 2008; Stagg et al., 2018). Also we suggest that as the medial prefrontal cortex calculates reward expectations in response to outcomes (Pagnoni et al., 2002), MFC inhibition by cathodal stimulation may have led to suppression of calculation of expected compensation, disrupting learning.

This result, however, should be considered together with the suppression of FRN in the cathodal stimulation group demonstrated earlier (Gorin et al., 2022), when, contrary to the prior hypothesis, the value of the component in the stimulation group was significantly higher than in the control group (Gorin et al., 2022). Interestingly, the ctDCS managed to make subjects more sensitive to differences in magnitude of presented feedback: their FRN of different monetary sums was different in cases when the difference was high (1 rub vs. 50 rub). To our knowledge, this is the first time when the cathodal tDCS of the MFC increased the performance monitoring process in such a way.

Our findings demonstrate the inconsistency of FRN and P2 amplitudes to this assumption, which we attribute to the multidirectional stimulation effect on higher-order neural networks. Such results could be due to the anatomical features of the structures responsible for decision-making: the sources of FRN are located in the dorsal ACC (Hauser et al., 2014). Although the main sources of P2 are in the anterior part of the auditory cortex, the lateral part of Heschl’s gyrus (Ross & Tremblay, 2009), and auditory association cortices (Crowley and Colrain, 2004), some researchers propose that nontemporal sources such as frontal areas can additionally contribute to its expression (Ferreira-Santos et al., 2012). Namely, new EEG-fMRI research suggested that dlPFC may also be a source of P2 (Mayhew et al., 2010). Because of the irregular distribution of tDCS, these cortical areas may be directly exposed to electrical current.

We suppose that the medial frontal cortex, which we stimulated, may overlap with one of the sources of P2, and fall under the influence of the electric current. This may also be due to the difficult predictability of tDCS-induced current propagation in an anisotropic environment (Datta et al., 2009) and the diverse impact of tDCS on various types of neurons located in different layers of cortex (Radman et al., 2009). Perhaps our results can add evidence to the arguments that P2 consists of subcomponents (García-Larrea et al., 1992; Silva et al., 2020). Whether this is correct, and whether we affected an early subcomponent of P2, which is the source of the observed plastic changes, remains to be studied in the future experiment, using the multichannel tDCS.

Since anodal and cathodal stimulation are associated with LTP- and LTD-type plastic changes (Pelletier & Cicchetti, 2015; Podda et al., 2016; Stagg et al., 2018), we suggest that in our study cathodal tDCS has inhibited P2 source and led to the suppression of LTP, which is probably responsible for the plastic changes in the potential that we observe in the control group. Our results may add evidence in favor of the fact that tES does affect plastic reorganization in the cerebral cortex due to changes not in the potential itself, but in its temporal dynamics.

Overall, the outcomes of various plasticity-inducing brain stimulation paradigms exhibit considerable variability (Wiethoff et al., 2014). Nevertheless, the extant literature offers limited insights into the response variability associated with transcranial direct current stimulation (tDCS). According to Medina et al. (2017), there may be a substantial “file-drawer” of unpublished negative results in the tDCS literature. The present study contributes to this nebulous landscape by elucidating the multidirectional effects of tDCS on the medial frontal cortex (MFC), thereby underscoring the imperative to devise more reliable protocols and enhance the comprehension of the individual and task-dependent determinants governing responsiveness.

### Limitations and perspectives

Initially, tDCS does not offer complete anatomical targeting (Datta et al., 2009). The heterogeneous propagation of the current may extend effects beyond the primary target, thereby distorting both data and interpretation. Given the intricate nature of high-order neuronal networks, we recommend employing imaging techniques to identify off-target effects. Despite these limitations, non-invasive brain stimulation techniques demonstrate significant potential.

In future studies, we propose utilizing other forms of electric and magnetic stimulation, including anodal tES, transcranial alternating current stimulation (tACS), and alternative montages in similar paradigms. For instance, researchers could focus specifically on the N1-P2 complex sources by placing the reference electrode on the temporal region and the active electrode as described in Reinhart and Woodman (2014). We advocate for using source analysis (via magnetoencephalography (MEG) or functional near-infrared spectroscopy (fNIRS)) to confirm or refute the suppression of the N1-P2 sources targeted by stimulation. Additionally, we emphasize the importance of employing equally populated cathodal, anodal, and sham stimulation groups to investigate the effects of such stimulation comprehensively, yielding more intricate results.

Regarding the paradigm, our study employed only negative reinforcement, where participants sought to avoid monetary losses. To obtain a more comprehensive perspective, future research should incorporate paradigms that include other types of reinforcement.

## Conclusion

The present work demonstrates that cathodal tDCS can influence both FRN and P2 components, highlighting the multidirectional effects of tES within the nonmotor domain. Since the study utilized a monetary task, we observed alterations in component magnitudes related to both auditory stimuli and feedback.

Contrary to our initial hypothesis, we found that FRN in the cathodal group increased, suggesting that plastic effects may have been enhanced by a stronger feedback signal. Meanwhile, the P2 amplitude remained unchanged, potentially indicating the inhibitory effect of tDCS on the P2 component. Our research provides further evidence for the unconventional cathodal effect of tDCS, characterized by increased, rather than decreased excitability for one component of the ERPs and no effect on the other, both obtained within the same experimental task – the monetary incentive delay (MID). Our observations underscore the need for future studies employing tDCS and encourage more cautious interpretation of tDCS-related effects in the absence of proper psychophysiological control.

## References

Antal, A., Terney, D., Poreisz, C., & Paulus, W. (2007). Towards unravelling task-related modulations of neuroplastic changes induced in the human motor cortex. European Journal of Neuroscience, 26(9), 2687–2691.

Arnott, S. R., Bardouille, T., Ross, B., & Alain, C. (2011). Neural generators underlying concurrent sound segregation. Brain research, 1387, 116–124.

Atienza, M., Cantero, J. L., & Dominguez-Marin, E. (2002). The time course of neural changes underlying auditory perceptual learning. Learning & Memory, 9(3), 138–150.

Balleine, B. W., Delgado, M. R., & Hikosaka, O. (2007). The role of the dorsal striatum in reward and decision-making. Journal of Neuroscience, 27(31), 8161–8165.

Batsikadze, G., Moliadze, V., Paulus, W., Kuo, M. F., & Nitsche, M. (2013). Partially non-linear stimulation intensity-dependent effects of direct current stimulation on motor cortex excitability in humans. The Journal of physiology, 591(7), 1987–2000.

Bear, M. F., & Malenka, R. C. (1994). Synaptic plasticity: LTP and LTD. Current opinion in neurobiology, 4(3), 389–399.

Been, G., Ngo, T. T., Miller, S. M., & Fitzgerald, P. B. (2007). The use of tDCS and CVS as methods of non-invasive brain stimulation. Brain research reviews, 56(2), 346–361.

Boroda, E., Sponheim, S. R., Fiecas, M., & Lim, K. O. (2020). Transcranial direct current stimulation (tDCS) elicits stimulus-specific enhancement of cortical plasticity. Neuroimage, 211, 116598.

Correia, M. J. (1998). Neuronal plasticity: adaptation and readaptation to the environment of space. Brain research reviews, 28(1-2), 61–65.

Crowley, K. E., and Colrain, I. M. (2004). A review of the evidence for P2 being an independent component process: age, sleep and modality. Clin. Neurophysiol. 115, 732–744.

Datta, A., Bansal, V., Diaz, J., Patel, J., Reato, D., & Bikson, M. (2009). Gyri-precise head model of transcranial direct current stimulation: improved spatial focality using a ring electrode versus conventional rectangular pad. Brain stimulation, 2(4), 201–207.

Davis, H., & Zerlin, S. (1966). Acoustic relations of the human vertex potential. The Journal of the Acoustical Society of America, 39(1), 109–116.

Delgado, M. R., Jou, R. L., & Phelps, E. A. (2011). Neural systems underlying aversive conditioning in humans with primary and secondary reinforcers. Frontiers in neuroscience, 5, 71.

Delgado, M. R., Labouliere, C. D., & Phelps, E. A. (2006). Fear of losing money? Aversive conditioning with secondary reinforcers. Social cognitive and affective neuroscience, 1(3), 250–259.

Delgado, M. R., Locke, H. M., Stenger, V. A., & Fiez, J. A. (2003). Dorsal striatum responses to reward and punishment: effects of valence and magnitude manipulations. Cognitive, Affective, & Behavioral Neuroscience, 3(1), 27–38.

Delhommeau, K., Micheyl, C., & Jouvent, R. (2005). Generalization of frequency discrimination learning across frequencies and ears: implications for underlying neural mechanisms in humans. Journal of the Association for Research in Otolaryngology, 6(2), 171–179.

Elias, G. A., Bieszczad, K. M., & Weinberger, N. M. (2015). Learning strategy refinement reverses early sensory cortical map expansion but not behavior: Support for a theory of directed cortical substrates of learning and memory. Neurobiology of learning and memory, 126, 39–55.

Fecteau, S., Knoch, D., Fregni, F., Sultani, N., Boggio, P., & Pascual-Leone, A. (2007). Diminishing risk-taking behavior by modulating activity in the prefrontal cortex: a direct current stimulation study. Journal of Neuroscience, 27(46), 12500–12505.

Ferreira-Santos, F., Silveira, C., Almeida, P. R., Palha, A., Barbosa, F., & Marques-Teixeira, J. (2012). The auditory P200 is both increased and reduced in schizophrenia? A meta-analytic dissociation of the effect for standard and target stimuli in the oddball task. Clinical Neurophysiology, 123(7), 1300–1308.

Ford, J. M., Roth, W. T., Menon, V., & Pfefferbaum, A. (1999). Failures of automatic and strategic processing in schizophrenia: comparisons of event-related brain potential and startle blink modification. Schizophrenia research, 37(2), 149–163.

Frank, M. J., & Claus, E. D. (2006). Anatomy of a decision: striato-orbitofrontal interactions in reinforcement learning, decision making, and reversal. Psychological review, 113(2), 300.

Galván, A. (2010). Neural plasticity of development and learning. Human brain mapping, 31(6), 879–890.

Ganguly, K., & Poo, M. M. (2013). Activity-dependent neural plasticity from bench to bedside. Neuron, 80(3), 729–741.

García-Larrea, L., Lukaszewicz, A.-C., & Mauguiére, F. (1992). Revisiting the oddball paradigm. Non-target vs neutral stimuli and the evaluation of ERP attentional effects. Neuropsychologia, 30(8), 723–741.

Garrido, M. I., Friston, K. J., Kiebel, S. J., Stephan, K. E., Baldeweg, T., & Kilner, J. M. (2008). The functional anatomy of the MMN: a DCM study of the roving paradigm. Neuroimage, 42(2), 936–944.

Godey, B., Schwartz, D., De Graaf, J. B., Chauvel, P., & Liegeois-Chauvel, C. (2001). Neuromagnetic source localization of auditory evoked fields and intracerebral evoked potentials: a comparison of data in the same patients. Clinical neurophysiology, 112(10), 1850–1859.

Gordon, P. C., Zrenner, C., Desideri, D., Belardinelli, P., Zrenner, B., Brunoni, A. R., & Ziemann, U. (2018). Modulation of cortical responses by transcranial direct current stimulation of dorsolateral prefrontal cortex: A resting-state EEG and TMS-EEG study. Brain stimulation, 11(5), 1024–1032.

Gorin A. et al. (2020) Cortical plasticity elicited by acoustically cued monetary losses: an ERP study. Scientific reports, 10(1), 1–14.

Gorin A. et al. (2022). Transcranial direct current electrostimulation modulates the component of the negativity of the result of the action in the monetary game. Journal of Higher Nervous Activity. I.P. Pavlov, 678–689.

Habibi, A., Wirantana, V., & Starr, A. (2014). Cortical activity during perception of musical rhythm: Comparing musicians and nonmusicians. Psychomusicology: Music, Mind, and Brain, 24(2), 125–135.

Hanley, C. J., Singh, K. D., & McGonigle, D. J. (2016). Transcranial modulation of brain oscillatory responses: A concurrent tDCS–MEG investigation. Neuroimage, 140, 20–32.

Hari, R., Hämäläinen, H., Hämäläinen, M., Kekoni, J., Sams, M., & Tiihonen, J. (1990). Separate finger representations at the human second somatosensory cortex. Neuroscience, 37(1), 245–249.

Hauser, T. U., Iannaccone, R., Stämpfli, P., Drechsler, R., Brandeis, D., Walitza, S., & Brem, S. (2014). The feedback-related negativity (FRN) revisited: new insights into the localization, meaning and network organization. Neuroimage, 84, 159–168.

Hayes, E. A., Warrier, C. M., Nicol, T. G., Zecker, S. G., & Kraus, N. (2003). Neural plasticity following auditory training in children with learning problems. Clinical neurophysiology, 114(4), 673–684.

Holroyd, C. B., Larsen, J. T., & Cohen, J. D. (2004). Context dependence of the event-related brain potential associated with reward and punishment. Psychophysiology, 41(2), 245–253.

Jacobson, L., Koslowsky, M., & Lavidor, M. (2012). tDCS polarity effects in motor and cognitive domains: a meta-analytical review. Experimental brain research, 216(1), 1–10.

Jamil, A., Batsikadze, G., Kuo, H. I., Labruna, L., Hasan, A., Paulus, W., & Nitsche, M. A. (2017). Systematic evaluation of the impact of stimulation intensity on neuroplastic aftereffects induced by transcranial direct current stimulation. The Journal of physiology, 595(4), 1273–1288.

Johnston, M. V. (2009). Plasticity in the developing brain: implications for rehabilitation. Developmental disabilities research reviews, 15(2), 94–101.

Kania, B. F., Wrońska, D., & Zięba, D. (2017). Introduction to neural plasticity mechanism. Journal of Behavioral and Brain Science, 7(2), 41–49.

Knechtel, L., Schall, U., Cooper, G., Ramadan, S., Stanwell, P., Jolly, T., & Thienel, R. (2014). Transcranial direct current stimulation of prefrontal cortex: an auditory event-related potential and proton magnetic resonance spectroscopy study. Neurology, Psychiatry and Brain Research, 20(4), 96–101.

Knight, R. T., Scabini, D., Woods, D. L., & Clayworth, C. (1988). The effects of lesions of superior temporal gyrus and inferior parietal lobe on temporal and vertex components of the human AEP. Electroencephalography and clinical Neurophysiology, 70(6), 499–509.

Knutson, B., Taylor, J., Kaufman, M., Peterson, R., & Glover, G. (2005). Distributed neural representation of expected value. Journal of Neuroscience, 25(19), 4806–4812.

Krigolson, O. E. (2018). Event-related brain potentials and the study of reward processing: Methodological considerations. International Journal of Psychophysiology, 132, 175–183.

Kringelbach, M. L. (2005). The human orbitofrontal cortex: linking reward to hedonic experience. Nature reviews neuroscience, 6(9), 691–702.

Krugliakova, E., Gorin, A., Fedele, T., Shtyrov, Y., Moiseeva, V., Klucharev, V., & Shestakova, A. (2019). The monetary incentive delay (MID) task induces changes in sensory processing: ERP evidence. Frontiers in Human Neuroscience, 382.

Krugliakova, E., Klucharev, V., Fedele, T., Gorin, A., Kuznetsova, A., & Shestakova, A. (2018). Correlation of cue-locked FRN and feedback-locked FRN in the auditory monetary incentive delay task. Experimental Brain Research, 236(1), 141–151.

Kühnis, J., Elmer, S., Meyer, M., and Jäncke, L. (2013). Musicianship boosts perceptual learning of pseudoword-chimeras: an electrophysiological approach. Brain Topogr. 26, 110–125.

Kunzelmann, K., Meier, L., Grieder, M., Morishima, Y., & Dierks, T. (2018). No effect of transcranial direct current stimulation of the auditory cortex on auditory-evoked potentials. Frontiers in neuroscience, 880.

Kuriki, S., Ohta, K., and Koyama, S. (2007). Persistent responsiveness of long-latency auditory cortical activities in response to repeated stimuli of musical timbre and vowel sounds. Cereb. Cortex 17, 2725–2732.

Leek, M. R., & Watson, C. S. (1984). Learning to detect auditory pattern components. The Journal of the Acoustical Society of America, 76(4), 1037–1044.

Lutz, K., & Widmer, M. (2014). What can the monetary incentive delay task tell us about the neural processing of reward and punishment. Neuroscience and Neuroeconomics, 3(3), 33–45.

Magnée, M. J., De Gelder, B., Van Engeland, H., & Kemner, C. (2008). Audiovisual speech integration in pervasive developmental disorder: evidence from event-related potentials. Journal of Child Psychology and Psychiatry, 49(9), 995–1000.

Malenka, R. C. (1995). REVIEW ■ : LTP and LTD: Dynamic and Interactive Processes of Synaptic Plasticity. The Neuroscientist, 1(1), 35–42.

Matsushita, R., Puschmann, S., Baillet, S., & Zatorre, R. J. (2021). Inhibitory effect of tDCS on auditory evoked response: simultaneous MEG-tDCS reveals causal role of right auditory cortex in pitch learning. Neuroimage, 233, 117915.

May, A., Hajak, G., Gänssbauer, S., Steffens, T., Langguth, B., Kleinjung, T., & Eichhammer, P. (2007). Structural brain alterations following 5 days of intervention: dynamic aspects of neuroplasticity. Cerebral cortex, 17(1), 205–210.

Mayhew, S. D., Dirckx, S. G., Niazy, R. K., Iannetti, G. D., & Wise, R. G. (2010). EEG signatures of auditory activity correlate with simultaneously recorded fMRI responses in humans. NeuroImage, 49(1), 849–864.

Monte-Silva, K., Kuo, M. F., Liebetanz, D., Paulus, W., & Nitsche, M. A. (2010). Shaping the optimal repetition interval for cathodal transcranial direct current stimulation (tDCS). Journal of neurophysiology, 103(4), 1735–1740.

Näätänen, R., & Picton, T. (1987). The N1 wave of the human electric and magnetic response to sound: a review and an analysis of the component structure. Psychophysiology, 24(4), 375–425.

Näätänen, R., Pakarinen, S., Rinne, T., & Takegata, R. (2004). The mismatch negativity (MMN): towards the optimal paradigm. Clinical neurophysiology, 115(1), 140–144.

Näätänen, R., Schröger, E., Karakas, S., Tervaniemi, M., & Paavilainen, P. (1993). Development of a memory trace for a complex sound in the human brain. Neuroreport: An International Journal for the Rapid Communication of Research in Neuroscience.

Nitsche, M. A., & Paulus, W. (2000). Excitability changes induced in the human motor cortex by weak transcranial direct current stimulation. The Journal of physiology, 527(Pt 3), 633.

Nitsche, M. A., & Paulus, W. (2001). Sustained excitability elevations induced by transcranial DC motor cortex stimulation in humans. Neurology, 57(10), 1899–1901.

Nitsche, M. A., Fricke, K., Henschke, U., Schlitterlau, A., Liebetanz, D., Lang, N., … & Paulus, W. (2003). Pharmacological modulation of cortical excitability shifts induced by transcranial direct current stimulation in humans. The Journal of physiology, 553(1), 293–301.

Nitsche, M. A., Jaussi, W., Liebetanz, D., Lang, N., Tergau, F., & Paulus, W. (2004). Consolidation of human motor cortical neuroplasticity by D-cycloserine. Neuropsychopharmacology, 29(8), 1573–1578.

Nitsche, M. A., Müller-Dahlhaus, F., Paulus, W., & Ziemann, U. (2012). The pharmacology of neuroplasticity induced by non-invasive brain stimulation: building models for the clinical use of CNS active drugs. The Journal of Physiology, 590(19), 4641–4662.

Nitsche, M. A., Schauenburg, A., Lang, N., Liebetanz, D., Exner, C., Paulus, W., & Tergau, F. (2003). Facilitation of implicit motor learning by weak transcranial direct current stimulation of the primary motor cortex in the human. Journal of cognitive neuroscience, 15(4), 619–626.

Nitsche, M. A., Seeber, A., Frommann, K., Klein, C. C., Rochford, C., Nitsche, M. S., … Tergau, F. (2005). Modulating parameters of excitability during and after transcranial direct current stimulation of the human motor cortex. The Journal of Physiology, 568(1), 291–303.

Noonan, M. P., Kolling, N., Walton, M. E., & Rushworth, M. F. S. (2012). Re-evaluating the role of the orbitofrontal cortex in reward and reinforcement. European Journal of Neuroscience, 35(7), 997–1010.

Orduña, I., Liu, E. H., Church, B. A., Eddins, A. C., and Mercado, E. (2012). Evoked-potential changes following discrimination learning involving complex sounds. Clin. Neurophysiol. 123, 711–719.

Pagnoni, G., Zink, C. F., Montague, P. R., & Berns, G. S. (2002). Activity in human ventral striatum locked to errors of reward prediction. Nature neuroscience, 5(2), 97–98.

Paiva, T. O., Almeida, P. R., Ferreira-Santos, F., Vieira, J. B., Silveira, C., Chaves, P. L., … & Marques-Teixeira, J. (2016). Similar sound intensity dependence of the N1 and P2 components of the auditory ERP: Averaged and single trial evidence. Clinical Neurophysiology, 127(1), 499–508.

Patil, I. (2021). Visualizations with statistical details: The ‘ggstatsplot’ approach. Journal of Open Source Software, 6(61), 3167, doi:10.21105/joss.03167.

Pelletier, S. J., & Cicchetti, F. (2015). Cellular and molecular mechanisms of action of transcranial direct current stimulation: evidence from in vitro and in vivo models. International Journal of Neuropsychopharmacology, 18(2).

Perrault, N., & Picton, T. W. (1984). Event-related potentials recorded from the scalp and nasopharynx. I. N1 and P2. Electroencephalography and Clinical Neurophysiology/Evoked Potentials Section, 59(3), 177–194.

Picton, T. (2013). Hearing in time: evoked potential studies of temporal processing. Ear and hearing, 34(4), 385–401.

Podda, M. V., Cocco, S., Mastrodonato, A., Fusco, S., Leone, L., Barbati, S. A., … & Grassi, C. (2016). Anodal transcranial direct current stimulation boosts synaptic plasticity and memory in mice via epigenetic regulation of Bdnf expression. Scientific reports, 6(1), 1–19.

Ponton, C. W., Vasama, J. P., Tremblay, K., Khosla, D., Kwong, B., & Don, M. (2001). Plasticity in the adult human central auditory system: evidence from late-onset profound unilateral deafness. Hearing research, 154(1-2), 32–44.

Purdy, S. C., Kelly, A. S., & Thorne, P. R. (2001). Auditory evoked potentials as measures of plasticity in humans. Audiology and Neurotology, 6(4), 211–215.

Radman, T., Ramos, R. L., Brumberg, J. C., & Bikson, M. (2009). Role of cortical cell type and morphology in subthreshold and suprathreshold uniform electric field stimulation in vitro. Brain stimulation, 2(4), 215–228.

Rauschecker, J. P. (1999). Auditory cortical plasticity: a comparison with other sensory systems. Trends in neurosciences, 22(2), 74–80.

Reinhart, R. M., & Woodman, G. F. (2014). Causal control of medial–frontal cortex governs electrophysiological and behavioral indices of performance monitoring and learning. Journal of Neuroscience, 34(12), 4214–4227.

Reinke, K. S., He, Y., Wang, C., & Alain, C. (2003). Perceptual learning modulates sensory evoked response during vowel segregation. Cognitive Brain Research, 17(3), 781–791.

Reis, J., Schambra, H. M., Cohen, L. G., Buch, E. R., Fritsch, B., Zarahn, E., … & Krakauer, J. W. (2009). Noninvasive cortical stimulation enhances motor skill acquisition over multiple days through an effect on consolidation. Proceedings of the National Academy of Sciences, 106(5), 1590–1595.

Ross, B., and Tremblay, K. (2009). Stimulus experience modifies auditory neuromagnetic responses in young and older listeners. Hear. Res. 248, 48–59.

Rouhani, N., & Niv, Y. (2021). Signed and unsigned reward prediction errors dynamically enhance learning and memory. Elife, 10.

Sambrook, T. D., & Goslin, J. (2014). Mediofrontal event-related potentials in response to positive, negative and unsigned prediction errors. Neuropsychologia, 61, 1–10.

Schultz, W. (2002). Getting formal with dopamine and reward. Neuron, 36(2), 241–263.

Seitz, A., and Watanabe, T. (2005). A unified model for perceptual learning. Trends Cogn. Sci. (Regul. Ed.) 9, 329–334.

Seppänen, M., Hämäläinen, J., Pesonen, A. K., and Tervaniemi, M. (2012). Music training enhances rapid neural plasticity of N1 and P2 source activation for unattended sounds. Front. Hum. Neurosci. 6:43.

Sheehan, K. A., Mcarthur, G. M., and Bishop, D. V. (2005). Is discrimination training necessary to cause changes in the P2 auditory event-related brain potential to speech sounds? Brain Res. Cogn. Brain Res. 25, 547–553.

Silva, D. M., Rothe-Neves, R., & Melges, D. B. (2020). Long-latency event-related responses to vowels: N1-P2 decomposition by two-step principal component analysis. International Journal of Psychophysiology, 148, 93–102.

Stagg, C. J., Antal, A., & Nitsche, M. A. (2018). Physiology of Transcranial Direct Current Stimulation. The Journal of ECT, 1.

Sutton, R. S., & Barto, A. G. (2018). Reinforcement learning: An introduction. MIT press.

Syka, J. (2002). Plastic changes in the central auditory system after hearing loss, restoration of function, and during learning. Physiological reviews, 82(3), 601–636.

Tadel, M. (2004). Gled-an Implementation of a hierarchic Server-client Model. Applied Parallel and Distributed Computing (Advances in Computation: Theory and Practice), 16.

Tang, M. F., & Hammond, G. R. (2013). Anodal transcranial direct current stimulation over auditory cortex degrades frequency discrimination by affecting temporal, but not place, coding. European Journal of Neuroscience, 38(5), 2802–2811.

Teyler, T. J., & DiScenna, P. (1987). Long-term potentiation. Annual review of neuroscience, 10(1), 131–161.

Tong, Y., Melara, R. D., and Rao, A. (2009). P2 enhancement from auditory discrimination training is associated with improved reaction times. Brain Res. 1297, 80–88.

Tremblay, K. L. (2003). Central auditory plasticity. The Hearing Journal, 56(1), 10.

Tremblay, K., Kraus, N., McGee, T., Ponton, C., & Otis, B. (2001). Central auditory plasticity: changes in the N1-P2 complex after speech-sound training. Ear and hearing, 22(2), 79–90.

Tsuchida, A., Doll, B. B., & Fellows, L. K. (2010). Beyond reversal: a critical role for human orbitofrontal cortex in flexible learning from probabilistic feedback. Journal of Neuroscience, 30(50), 16868–16875.

Vitor-Costa, M., Okuno, N. M., Bortolotti, H., Bertollo, M., Boggio, P. S., Fregni, F., & Altimari, L. R. (2015). Improving cycling performance: transcranial direct current stimulation increases time to exhaustion in cycling. PloS one, 10(12), e0144916.

Walsh, M. M., & Anderson, J. R. (2012). Learning from experience: event-related potential correlates of reward processing, neural adaptation, and behavioral choice. Neuroscience & Biobehavioral Reviews, 36(8), 1870–1884.

Weinberger, N. M. (1995). Dynamic Regulation of Receptive Fields and Maps in the Adult Sensory Cortex. Annual Review of Neuroscience, 18(1), 129–158.

Weinberger, N. M., & Diamond, D. M. (1987). Physiological plasticity in auditory cortex: rapid induction by learning. Progress in neurobiology, 29(1), 1–55.

Wilkinson, R. T., & Lee, M. V. (1972). Auditory evoked potentials and selective attention. Electroencephalography and Clinical Neurophysiology, 33(4), 411–418.

Wisniewski, M. G., Ball, N. J., Zakrzewski, A. C., Iyer, N., Thompson, E. R., & Spencer, N. (2020). Auditory detection learning is accompanied by plasticity in the auditory evoked potential. Neuroscience Letters, 134781.

Woods, A. J., Antal, A., Bikson, M., Boggio, P. S., Brunoni, A. R., Celnik, P., … & Nitsche, M. A. (2016). A technical guide to tDCS, and related non-invasive brain stimulation tools. Clinical neurophysiology, 127(2), 1031–1048.

Xiong, X. R., Liang, F., Zingg, B., Ji, X. Y., Ibrahim, L. A., Tao, H. W., & Zhang, L. I. (2015). Auditory cortex controls sound-driven innate defense behaviour through corticofugal projections to inferior colliculus. Nature communications, 6(1), 1–12.

Zaehle, T., Beretta, M., Jäncke, L., Herrmann, C. S., & Sandmann, P. (2011). Excitability changes induced in the human auditory cortex by transcranial direct current stimulation: direct electrophysiological evidence. Experimental brain research, 215(2), 135–140.

Zwislocki, J., Maire, F., Feldman, A. S., & Rubin, H. (1958). On the effect of practice and motivation on the threshold of audibility. The Journal of the Acoustical Society of America, 30(4), 254–262.

